# The Y chromosome gene *draupnir* reveals constraints on engineering Y-linked sex-ratio distorters in malaria mosquitoes

**DOI:** 10.64898/2026.04.03.716343

**Authors:** Rocco D’Amato, Elad Shmuel Yonah, Alessia Cagnetti, Flavia Krsticevic, Arad Sarig, Salvatore Di Martino, Alessandro Trusso, Roberto Galizi, Nikolai Windbichler, Alekos Simoni, Philippos Aris Papathanos

**Author notes:** These authors contributed equally to this work (Rocco D’Amato, Elad Shmuel Yonah, Alessia Cagnetti).

## Abstract

Y chromosome sex distorters offer a powerful strategy for mosquito population suppression, but their implementation is constrained by the transcriptional silencing of Y-linked transgenes during spermatogenesis. To investigate how endogenous Y genes evade this repression, we characterized *draupnir* (formerly *YG5*), a multicopy Y-linked gene of *Anopheles gambiae* encoding a Zip3-like meiotic protein. Comparative genomic and phylogenetic analyses revealed that *draupnir* originated through duplication of the autosomal paralog *skirnir*, itself derived from the ancestral recombination factor *vilya*, followed by amplification into a tandem array on the Y chromosome. We show that *draupnir* is actively transcribed during meiosis, making it the only known Y-linked gene expressed during mosquito spermatogenesis.

To test whether its regulatory region is sufficient to confer sperm-specific expression from the Y chromosome, we cloned the *draupnir* promoter to drive expression of an X-chromosome shredder. When inserted on an autosome, the construct drove meiotic expression and strong sex-ratio distortion in the progeny of transgenic males. In contrast, the same construct inserted on the Y chromosome was transcriptionally silent and produced balanced sex ratios. These results demonstrate that *draupnir* expression depends on its native genomic context within a multicopy Y-linked array and that its promoter alone cannot overcome transcription silencing. Our findings reveal a fundamental constraint on Y-chromosome-based genetic control strategies and point to future approaches for enabling transcription from otherwise repressed sex chromosomes.

## Introduction

Y chromosomes are transmitted exclusively from fathers to sons. Their evolution is sculpted by this isolation through selective forces acting solely in the interests of male fitness. This property can be exploited for genetic control using engineered sex ratio distorters (SRDs) located on the Y, designed to suppress or even eradicate pest populations by targeting and cleaving X-chromosome linked sequences during male meiosis, eliminating X-bearing gametes and producing an excess of male offspring (Windbichler, Papathanos and Crisanti, 2008; Galizi *et al*., 2014, 2016; Fasulo *et al*., 2020; Haber *et al*., 2024). Because such Y-drives are never present in females, they are shielded from counter-selection and can achieve high efficacy and long-term stability once introduced into the wild, according to modelling (Deredec, Godfray and Burt, 2011; Burt and Deredec, 2018). Autosomal SRDs have demonstrated the feasibility of this approach in the lab, but these are limited by Mendelian inheritance and negative selection in females.

A major obstacle to implementing Y-drives in *Anopheles gambiae* (*An. gambiae*) mosquitoes is the transcriptional silencing of the Y chromosome during male meiosis. Like other species with heteromorphic sex chromosomes, *An. gambiae* exhibits meiotic sex-chromosome inactivation (MSCI), in which both X and Y chromosomes are transcriptionally repressed from early pachytene onward (Taxiarchi *et al*., 2019; Page *et al*., 2023). As a result, transgenes driven by canonical meiotic promoters such as *β2-tubulin* fail to express when inserted on the Y chromosome (Alcalay *et al*., 2021), effectively preventing implementation of Y-linked SRDs. Nevertheless, in many species, Y-linked genes have evolved testis-specific expression apparently as an adaptation to overcome MSCI. In *Drosophila melanogaster*, for instance, Y-linked genes such as *kl-2*, *kl-3*, and *kl-5* are expressed predominantly in late spermatogenesis (Bonaccorsi *et al*., 1988; Vibranovski, Koerich and Carvalho, 2008; Carvalho, Koerich and Clark, 2009; Zhang *et al*., 2020). Similar patterns occur in mammals, where numerous ampliconic Y genes, such as *Sly*, *Ssty*, and *Zfy*, show testis- and sperm-specific expression (Cocquet *et al*., 2009; Riel *et al*., 2013; Comptour *et al*., 2014; Rathje *et al*., 2019; Subrini and Turner, 2021).

We previously identified an *An. gambiae* Y-linked gene, *YG5*, with sperm-specific expression (Hall *et al*., 2016). In that study, we reported that *YG5* likely originated by duplication of the autosomal gene AGAP013757, including ∼2 kb of upstream flanking DNA. We showed that *YG5* is expressed exclusively in testes, together with its autosomal paralog, and that it is physically located at one tip of the mitotic Y chromosome within a region enriched in the Y-specific transposable element *changuu*. Read-depth analysis suggested that *YG5* is present in multiple Y-linked copies, approximately tenfold higher than a single-copy locus in males, and that its copy number is remarkably stable across wild-type Y chromosomes from field-collected individuals in Cameroon, unlike other Y-chromosome loci. However, the biological role and developmental timing of *YG5* expression remained unresolved. Here, we revisit *YG5* to define its evolutionary origin, genomic organization, and expression dynamics, and to directly test whether its regulatory sequences can overcome transcriptional suppression to support functional Y-linked SRDs.

## Results

### The Y chromosome gene *YG5* encodes a Zip3-like meiotic protein

Using tBLASTn, we identified a second *YG5* paralog in the *An. gambiae* genome, AGAP006749 located on chromosome 2L, in addition to the previously known paralog AGAP013757 located on chromosome 3R (Hall *et al*., 2016). Including AGAP006749 in homology searches with *D. melanogaster* revealed three structurally-related Zip3-like genes*: vilya* (CG2709), *nenya* (CG31053), and *narya* (CG12200). First characterized in yeast, Zip3-like proteins function as pro-crossover factors during meiosis, localizing along the synaptonemal complex (Ouspenski, Elledge and Brinkley, 1999; Agarwal and Roeder, 2000). In *D. melanogaster*, the Zip3-like proteins *vilya*, *nenya* and *narya* facilitate the initiation of recombination during meiotic prophase and are essential for the proper formation of meiotic double-stranded DNA breaks during early pachytene (Lake *et al*., 2015, 2019). Mutations in *vilya* cause high levels of nondisjunction at the first meiotic division in females, while the combined absence of both *nenya* and *narya* abolishes recombination, leading to complete segregation failure. Interestingly, the presence of multiple Zip3-like genes within an organism is common and, in some cases, paralogs perform partially redundant roles (Zhang *et al*., 2018; Lake *et al*., 2019).

To better understand the evolutionary relationships between the mosquito and fly genes, we conducted a comparative analysis of protein sequences, gene structures and genomic synteny. The results indicate that AGAP006749 represents the true ortholog of *vilya*, whereas the remaining pairs arose from *vilya* independently in mosquitoes and flies (**Figure 1**). While *vilya* and *nenya* are present across the *Drosophila* genus, *narya* arose as a gene duplication event of *nenya* around 40 Mya, prior to the separation of the *melanogaster* subgroup (Lake *et al*., 2019). Like *nenya* and *narya*, *YG5* and AGAP013757 are intronless, whereas *vilya* and AGAP006749 contain introns (**Figure 1**).

**Figure 1:**
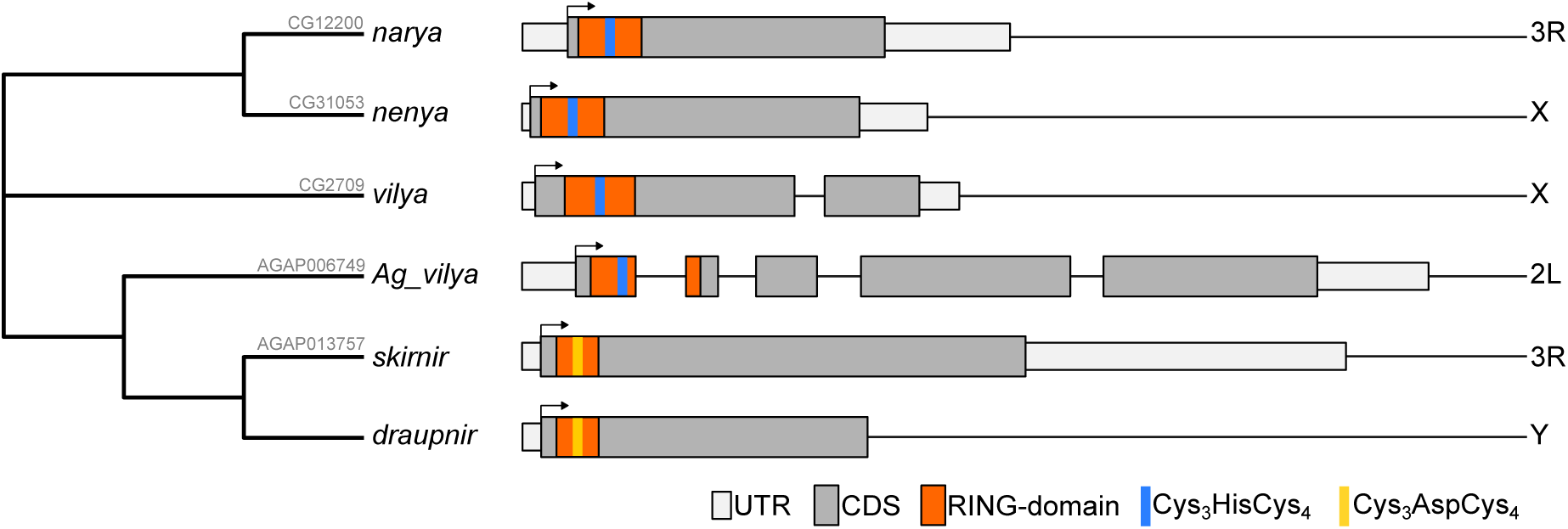
Gene structure and chromosomal organization of Zip3-like genes in *Drosophila melanogaster* and *Anopheles gambiae*. Cladogram (left) showing the inferred evolutionary relationships among the three *D. melanogaster* Zip3-like paralogs (*narya*, *nenya*, *vilya*) and their *An. gambiae* counterparts (*Ag_vilya*, *skirnir*, *draupnir*), with fly gene identifiers (CG numbers) and mosquito gene identifiers (AGAP numbers) indicated. Gene structure diagrams (right) show the organization of each locus. Boxes represent UTR (white), CDS (gray), and RING-finger domain (red); Cys_3_HisCys_4_ and Cys_3_AspCys_4_ zinc-coordinating motifs are highlighted in blue and orange, respectively.

Like all Zip3-like proteins, *vilya*, *nenya* and *narya* contain a RING finger domain in their N-terminus composed of the Cys_3_HisCys_4_ motif, which coordinates two zinc ions and is critical for their E3 SUMO ligase activity (Deshaies and Joazeiro, 2009). This motif is conserved in AGAP006749 but modified in both *YG5* and AGAP013757, where the histidine residue is replaced by an aspartic acid, further supporting their independent evolutionary origin. The RING domain and their existence as a functional triplet in *D. melanogaster* motivated their naming after Tolkien’s three elven rings of power (*vilya*, *nenya* and *narya*). To continue with this theme, we have renamed *YG5* as *draupnir*, after Odin’s self-replicating ring and AGAP013757 as *skirnir*, after the messenger who delivered *draupnir*. We refer to AGAP006749 as *Ag_vilya* when comparing across species, or *vilya* for brevity.

### Evolution of Zip3-like genes in mosquitoes

To investigate the evolutionary history of the Zip3-like genes across mosquitoes, we next searched for homologs of *Ag_vilya*, *skirnir* and *draupnir* in representative genome assemblies spanning the *Anopheles* phylogeny, as well as *Aedes aegypti*, *Culex quinquefasciatus* and *D. melanogaster*. After manually curating all predicted coding sequences to remove sequencing or annotation errors, we identified 35 homologs and used these to construct a maximum-likelihood phylogenetic tree based on protein sequences (**Figure 2**). The resulting tree reveals a stepwise expansion of Zip3-like genes within *Anopheles*. The ancestral *vilya* ortholog is found across *Culex*, *Aedes*, and all *Anopheles* subgenera, including *Cellia*, *Anopheles sensu stricto,* and *Nyssorhynchus*, indicating that it predates the divergence between *Anophelinae* and *Culicinae* (>200 Mya) and represents the ancestral single-copy *Zip3*-like gene in mosquitoes. Within *Anopheles*, a duplication of *vilya* gave rise to the autosomal paralog *skirnir*, which is present throughout the genus but absent from *Culicines*. The broad distribution of *skirnir* orthologs across *Cellia*, *Anopheles s. s.,* and *Nyssorhynchus* lineages implies that this duplication occurred early in *Anopheles* evolution, before the diversification of these subgenera, approximately 100-120 Mya. A subsequent duplication gave rise to *draupnir*, which we detected in *An. gambiae*, *An*. *coluzzii*, and *An*. *melas*. We did not find clear *draupnir* orthologs in other members of the *An. gambiae* complex (e.g. *An. arabiensis, An. merus*), but given the incompleteness of Y-chromosome assemblies, this absence may reflect lineage loss, assembly gaps, or reduced copy number below detection, rather than true absence. Accordingly, we interpret *draupnir* as restricted to, or at least enriched in, the *An. gambiae* complex, pending targeted Y-linked sequencing in additional complex members.

**Figure 2:**
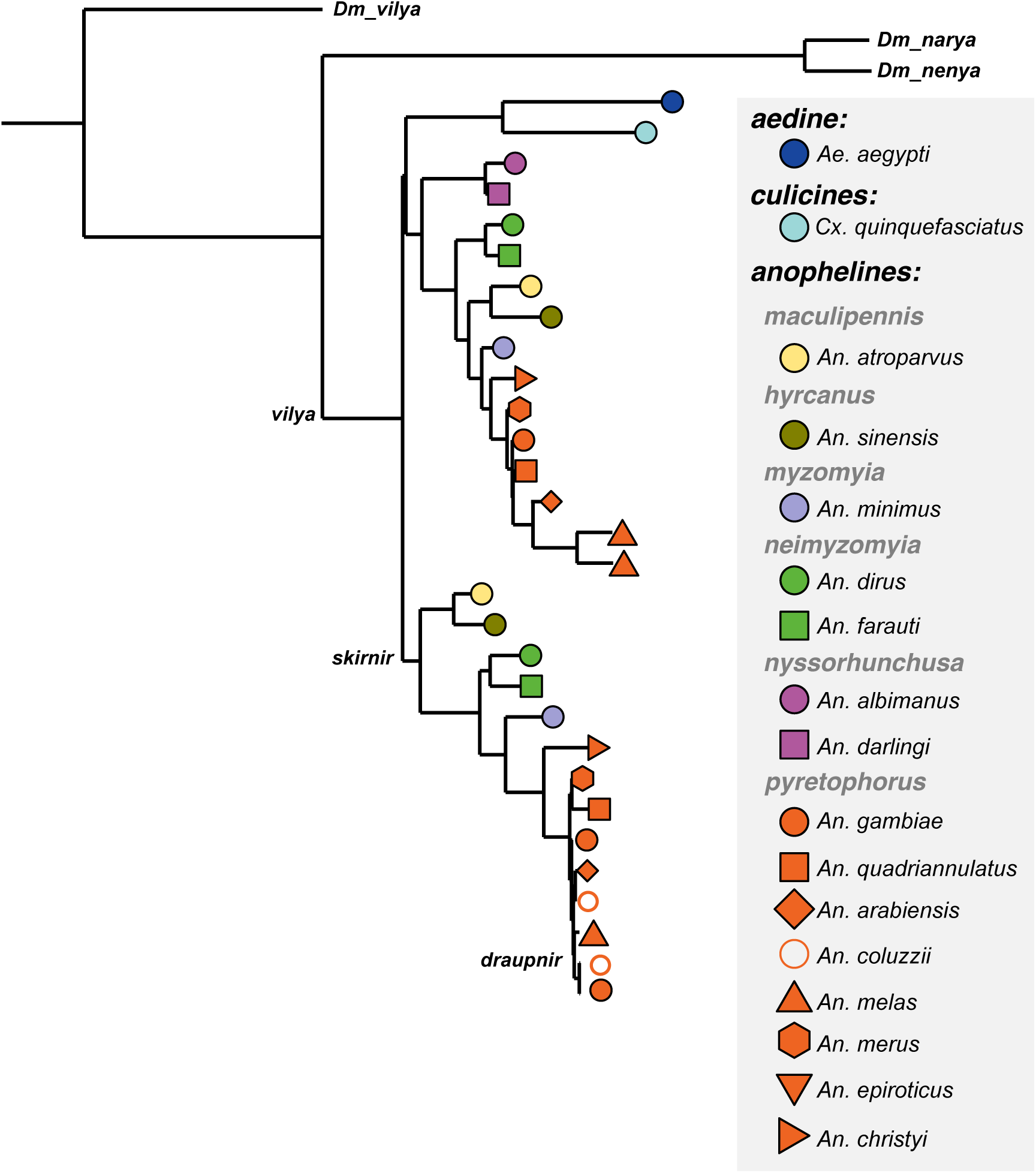
Phylogenetic distribution of Zip3-like genes across mosquitoes. Maximum-likelihood phylogenetic tree of Zip3-like protein sequences from *D.* melanogaster (outgroup), *Aedes aegypti*, *Culex quinquefasciatus*, and representative *Anopheles* species spanning the major subgenera. Three mosquito clades are indicated: *vilya* (ancestral ortholog, present in all species), *skirnir* (*Anopheles*-restricted autosomal paralog), and *draupnir* (restricted to the *An. gambiae* complex). Anopheline subgeneric groupings are labeled in gray italic text. Species are represented by colored shapes as shown in the key. *D. melanogaster* sequences (*vilya*, *narya*, *nenya*) are included as reference.

### Organization of the *draupnir* array on the Y chromosome

To resolve the structure of the *draupnir* locus, we screened a previously generated Bacterial Artificial Chromosome (BAC) library constructed from male genomic DNA of the *An. gambiae* G3 strain (Bernardini *et al*., 2014). PCR screening using *draupnir*-specific primers identified two positive clones, BAC-2C and BAC-5B, which were subsequently sequenced using PacBio reads. Reads spanning the complete BACs were selected for analysis to avoid assembler-associated artefacts generated by repetitive sequences. To confirm that both BACs originated from the Y chromosome, male and female whole-genome sequencing illumina libraries from the *An. gambiae* G3 strain were mapped to the BACs using Bowtie (Langmead *et al*., 2009). Normalized coverage indicated strong enrichment of male reads across both BACs, with negligible female coverage in the region spanning the *draupnir* array (**Figure 3A-C**), confirming their Y-linked origin. Annotation of the long reads using known *An. gambiae* Y-sequences identified multiple full-length *draupnir* open reading frames (ORFs) arranged in a head-to-tail tandem array. BAC-2C contained nine consecutive *draupnir* copies (*draupnir* 1-9) (**Figure 3A**), while BAC-5B contained a single *draupnir* coding sequence (*draupnir* 10) (**Figure 3C**). Each *draupnir* copy was separated by intergenic regions enriched for the Y-specific transposable element *changuu*, consistent with previous data of linkage between the two sequences (Hall *et al*., 2016). The distances between consecutive *draupnir* copies varied from approximately 13.6 kb to 23.2 kb, suggesting local heterogeneity in the repeat structure and incomplete homogenization among array units. Alignment of all ten Y-linked *draupnir* ORFs revealed near-complete sequence identity across the array (**Figure 3D**). Among the nine BAC-2C copies, *draupnir-5* carried a single SNP, while *draupnir-10* on BAC-5B harboured three SNPs and is truncated by 32 bp at the 3’ end; all substitutions resulted in non-synonymous changes to the predicted protein sequence. In contrast, alignment with the autosomal paralog *skirnir* revealed 27 divergent positions and an ORF 318 bp longer than any Y-linked copy, indicating substantially greater sequence divergence between the Y-linked array and its autosomal ancestor. These findings strongly support a single ancestral duplication event in which *skirnir* moved to the Y chromosome, followed by local amplification to generate the multicopy *draupnir* array and subsequent sequence homogenization among array units.

**Figure 3:**
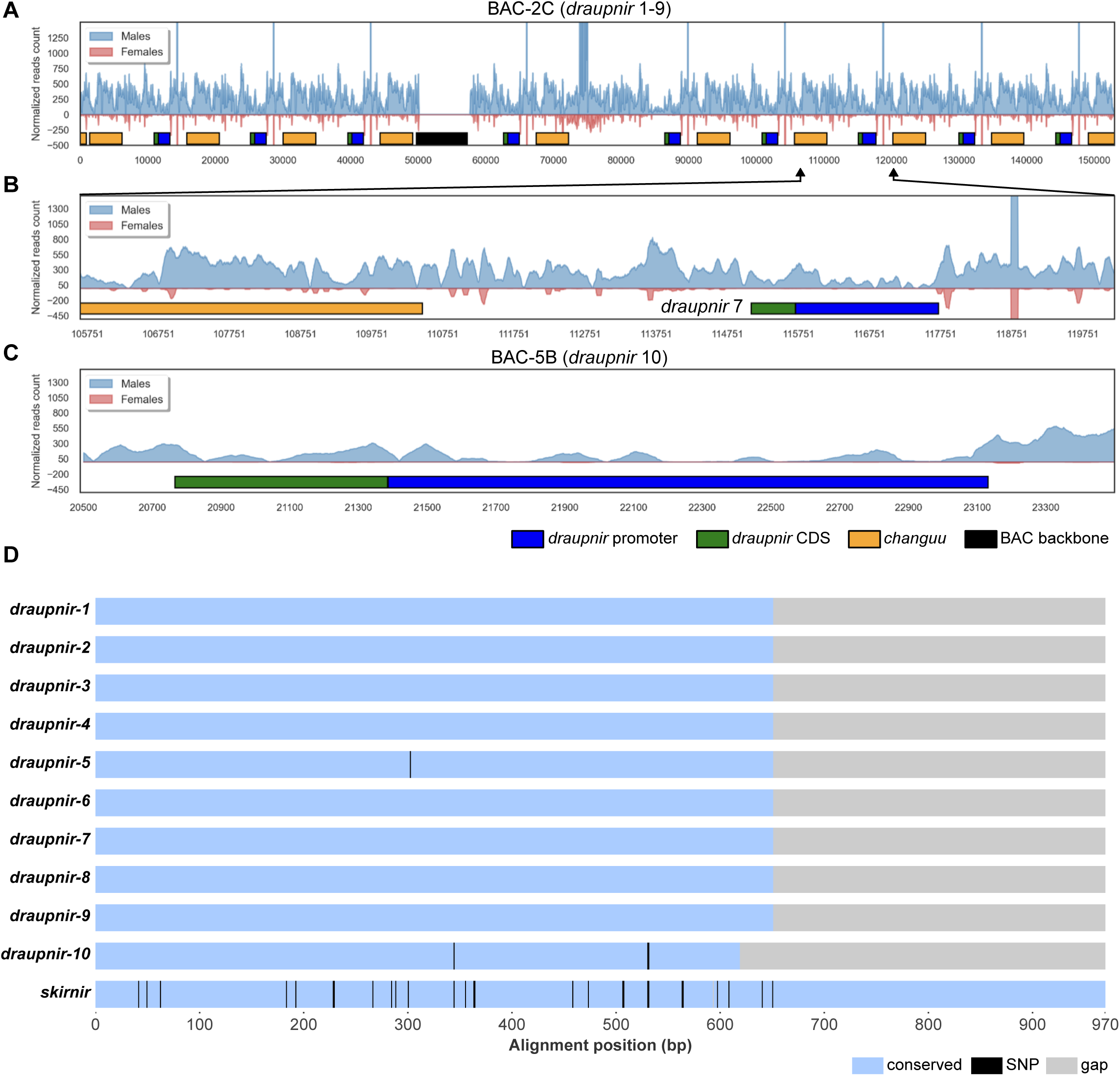
Genomic organization and Y-linked origin of the *draupnir* locus. Normalized DNA sequencing coverage from male (blue) and female (pink) whole-genome Illumina libraries mapped to the two *draupnir*-positive BAC clones. Coverage values for female reads are plotted inverted (below zero) for clarity. Annotated features are shown as colored bars below each coverage track: *draupnir* upstream regulatory regions (blue), *draupnir* coding sequences (green), *changuu* transposable elements (orange), and the pIndigoBAC5 vector backbone (black). (**A**) Full-length coverage of BAC-2C (∼150 kb). Strong and uniform enrichment of male reads with negligible female coverage across the *draupnir*-containing array confirms Y-chromosome origin. Nine *draupnir* copies are arranged in a head-to-tail tandem array, each separated by *changuu*-enriched intergenic intervals. (**B**) Zoomed view of a representative single *draupnir* repeat unit within BAC-2C (coordinates ∼ 105.7–119.7 kb), illustrating the typical spacing between consecutive copies and the relative sizes of the upstream regulatory region, coding sequence, and flanking *changuu* elements. (**C**) Full-length coverage of BAC-5B, which contains the single *draupnir*-10 ORF. Male-specific coverage confirms its Y-linked origin. The single copy spans a region of ∼2.5 kb, consistent with the structure of individual units in the BAC-2C array. **(D)** Pairwise alignment of the ten Y-linked *draupnir* ORFs and the autosomal paralogue *skirnir*. Each row represents one sequence; conserved positions are shown in blue, single nucleotide polymorphisms (SNPs) in black, and alignment gaps in grey. *draupnir*-5 carries a single SNP relative to all other Y-linked copies; *draupnir*-10 harbours three SNPs and is truncated by 32 bp at its 3’ end. *skirnir* has 27 divergent positions with the *draupnir* consensus and encodes an ORF 318 bp longer, reflecting its greater divergence as the autosomal ancestral copy.

### Expression of *draupnir* during spermatogenesis

To confirm and refine the tissue and developmental specificity of *draupnir* expression, we designed primers targeting diagnostic SNPs that distinguish *draupnir* from its autosomal paralog *skirnir*. RT-PCR amplification produced a single product exclusively in testes, with no detectable expression in male accessory glands (MAGs), female heads, carcasses, or ovaries (**Figure 4A**). Sequencing of the amplicon verified the presence of SNPs characteristic of *draupnir* (**Figure 4B**). Quantitative RT-PCR further showed that testis expression was approximately 100-fold higher than in any other tissue (**Figure 4C**), confirming strong expression during spermatogenesis, consistent with earlier observations that *skirnir* and its Y-linked duplicate are testis-expressed (Hall *et al*., 2016).

**Figure 4:**
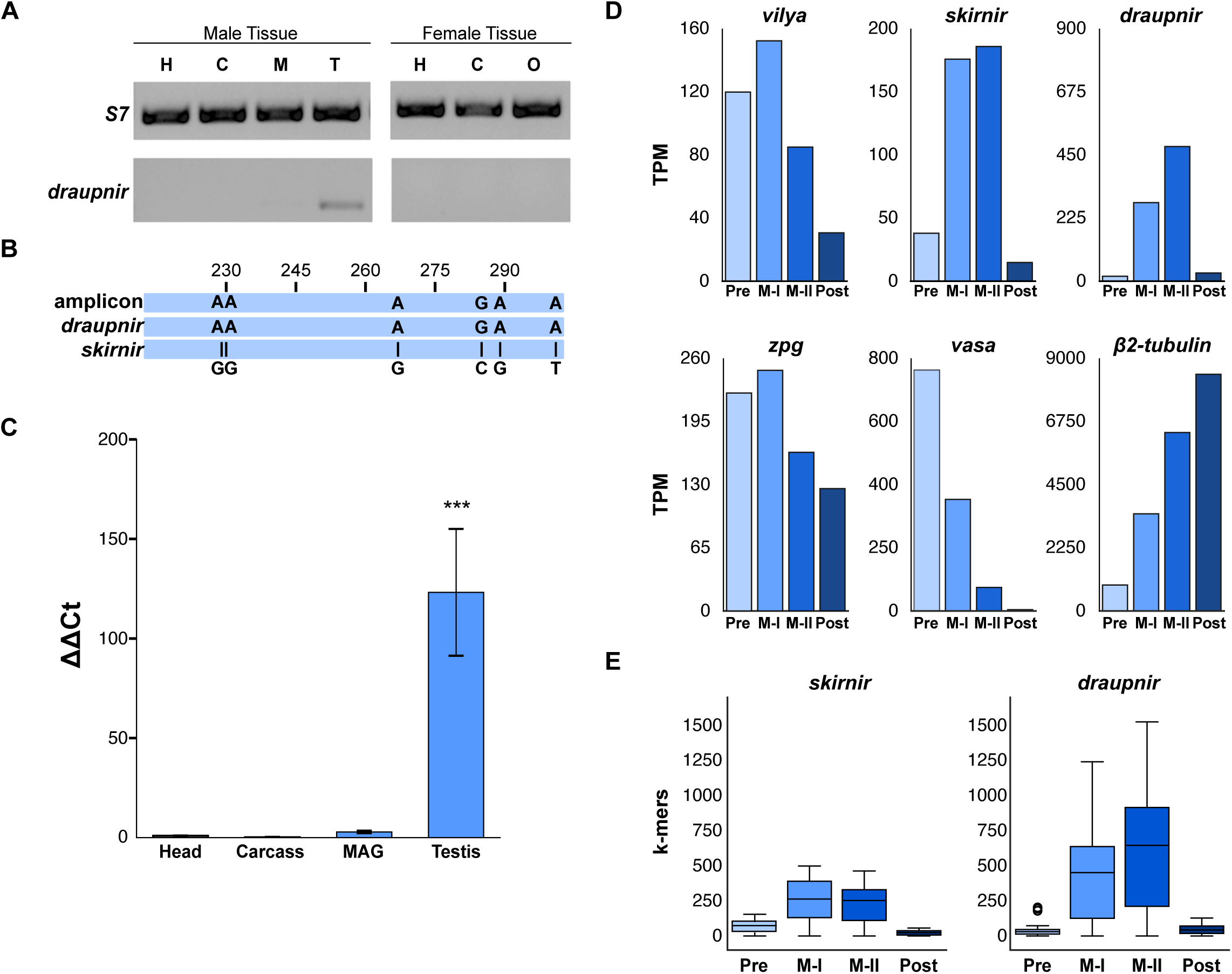
Expression of *draupnir* is testis-specific and peaks during meiosis. **(A)** Qualitative RT-PCR analysis of *draupnir* expression across dissected male (H, head; C, carcass; M, MAGs; T, testes) and female (H, head; C, carcass; O, ovaries) tissues from wild-type *An. gambiae* G3 mosquitoes. The ribosomal protein gene *S7* served as a loading control. A single *draupnir* amplicon is detected exclusively in testes. **(B)** Nucleotide alignment of the testis cDNA amplicon against the genomic sequences of *draupnir* and *skirnir*. The amplicon sequence is identical to *draupnir* at all positions; diagnostic substitutions distinguishing *draupnir* from *skirnir* (indicated below) confirm that the testis-expressed transcript derives from *draupnir* and not from its autosomal paralog. **(C)** Quantitative RT-PCR showing ΔΔCt values for *draupnir* expression normalized to *Rpl19* and calibrated to head tissue. Statistical comparisons were performed by one-way ANOVA followed by Dunnett’s post-hoc test against head tissue (ΔCt values, *n* = 3 biological replicates). \*\*\**p* < 0.001. **(D)** Stage-resolved RNA-seq expression profiles (TPM) of *vilya*, *skirnir*, *draupnir*, and the reference genes *zpg*, *vasa*, and *β2-tubulin* across four spermatogenic stages: pre-meiotic (Pre), meiosis I (M-I), meiosis II (M-II), and post-meiotic (Post), from Taxiarchi et al. (2019). **(E)** k-mer–based quantification of paralog-specific transcripts for *skirnir* and *draupnir* across spermatogenic stages. Box plots show the distribution of k-mer counts per stage; *draupnir*-specific k-mers are consistently more abundant than those of *skirnir* at all stages.

To determine the temporal dynamics of *draupnir* expression during spermatogenesis, we re-analyzed the stage-resolved RNA-seq dataset from Taxiarchi et al. 2019, which profiled four stages of spermatogenesis: pre-meiotic, meiotic (primary and secondary spermatocytes), and post-meiotic cells (Taxiarchi *et al*., 2019). Reads from each stage were mapped to *draupnir*, *skirnir* and the ancestral *vilya*. For comparison, we included *zpg* and *vasa* as pre-meiotic markers and *β2-tubulin* as a canonical meiotic marker. Expression of *vilya* began prior to meiosis and peaked in early spermatocytes, whereas both *draupnir* and *skirnir* were most highly expressed in secondary spermatocytes (**Figure 4D**). As expected, *zpg* and *vasa* were expressed from the earliest stages, while *β2-tubulin* expression increased progressively throughout spermatogenesis peaking in post-meiotic spermatids.

Given the high nucleotide identity (∼96%) between *draupnir* and *skirnir*, we employed a *k-mer*–based approach to distinguish paralog-specific expression. Using Jellyfish (Marçais and Kingsford, 2011), we generated all possible 25-*mer* sequences from each gene and identified those unique to any of the ten BAC-derived *draupnir* copies, those unique to *skirnir*, or shared between both. Corresponding *k-mers* were then extracted from the staged RNA-seq reads of Taxiarchi et al. (2019) and intersected with these diagnostic sets to quantify coverage across each spermatogenic stage. *Draupnir-*specific *k-mers* were consistently more abundant than those of *skirnir* at all stages, confirming that *draupnir* transcripts are the primary source of meiotic expression (**Figure 4E**). Thus, autosomal *skirnir* contributes minimally to the observed meiotic transcription signal, indicating that *draupnir* is the predominant source of expression during meiosis.

### Cloning and functional testing of the *draupnir* promoter

Since *draupnir* is expressed during meiosis, we hypothesized that its regulatory regions may have evolved mechanisms to sustain transcription despite MSCI. To test this, we designed primers to specifically amplify the *draupnir* 5’ regulatory regions (hereafter referred to as the promoter), leveraging sequence differences ∼ 2 kb upstream of the start codon that are conserved across all *draupnir* copies, but absent from *skirnir*. A single amplicon of the expected size (1989 bp) was amplified from male but not female genomic DNA and contained all *draupnir*-specific SNPs (**Supplementary Figure S1**). This putative promoter was cloned upstream of an I-*Ppo*I X-shredder cassette based on the destabilized I-PpoI*^124L^* variant fused to eGFP with the F2A ribosome skipping site and terminated by the *β2-tubulin* 3’ UTR (Galizi *et al*., 2014) (**Figure 5A**). The construct also included an attB site upstream of a promoterless eCFP marker for site-specific integration in docking strains, in which the attP site is located between the 3xP3 promoter and the DsRed marker, resulting in marker switching (Bernardini *et al*., 2014) (**Figure 5A**). Eggs from two docking strains were injected: one located on the Y chromosome (Y-attP (Bernardini *et al*., 2014)) and one on chromosome 2R (AttP-RFP-H2). Transgenic F1 larvae expressing eCFP were reared to adulthood and backcrossed to wild type individuals to establish stable strains, designated Draup-A and Draup-Y. As expected, transgenic larvae eclosed exclusively as males when the construct was inserted on the Y-chromosome, whereas both sexes were recovered when inserted on the autosome. From F2 onward, both transgenic strains were maintained by backcrossing to wild-type individuals. Draup-A was propagated through females, whereas Draup-Y was maintained through transgenic males.

**Figure 5:**
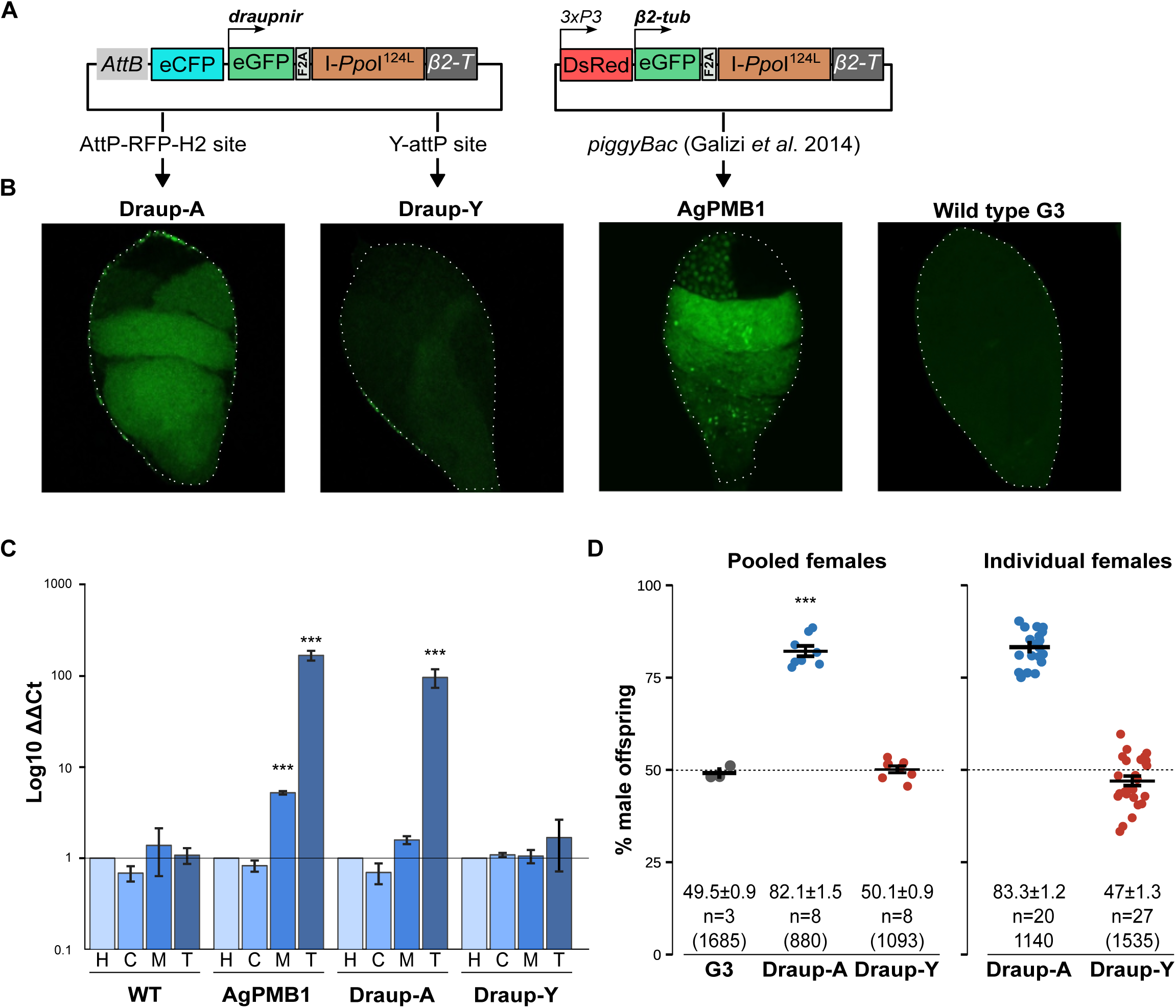
The *draupnir* promoter drives meiotic X-shredder expression from an autosomal but not Y-linked insertion. (**A**) Schematic of the transgenic constructs used to generate Draup-A (autosomal, chromosome 2R AttP-RFP-H2 docking site) and Draup-Y (Y-linked, Y-attP docking site). Both constructs carry the *draupnir* promoter driving a destabilized eGFP-I-*Ppo*I*^124L^*fusion via an F2A ribosome-skipping site, terminated by the *β2-tubulin* 3’ UTR, with an upstream promoterless eCFP marker flanked by attB for *φC31*-mediated site-specific integration. The AgPMB1 control strain (Galizi et al., 2014) uses a *piggyBac* insertion with a *β2-tubulin* promoter driving the same eGFP-I-*Ppo*I*^124L^* cassette. (**B**) Representative confocal fluorescence images of dissected testes from adult Draup-A, Draup-Y, AgPMB1, and wild-type G3 males, showing eGFP expression. Draup-A testes exhibit weak sperm eGFP signal; Draup-Y and wild-type testes show no detectable fluorescence; AgPMB1 shows robust expression. Dashed outlines indicate testes boundaries. (**C**) Quantitative RT-PCR of I-*Ppo*I transcript levels (Log₁₀ ΔΔCt, normalized to *Rpl19* and calibrated to head) across male tissues (H, head; C, carcass; M, MAGs; T, testes) from WT, AgPMB1, Draup-A, and Draup-Y adult males. Statistical comparisons for each strain were performed by one-way ANOVA followed by Dunnett’s post-hoc test against head tissue of each strain (ΔCt values, n = 2 biological replicates). ***p < 0.001. (**D**) Percentage of male offspring produced by Draup-A and Draup-Y males crossed to wild-type females, assessed as pooled (*en masse*, left) and individual female (right) experiments. Each dot represents one cage or one female. Horizontal bars show mean ± SEM; n indicates the number of replicates and total number of offspring sexed is indicated below each group. A dashed line marks the 50% expected sex ratio.

To test whether the *draupnir* promoter drives expression during spermatogenesis, we dissected the reproductive tissues from adult Draup-A, Draup-Y, AgPMB1 (autosomal *β2-tubulin*–driven I-PpoI*^124L^* X-shredder; (Galizi *et al*., 2014)), and wild-type males. Draup-Y testes showed no detectable eGFP fluorescence, whereas Draup-A testes exhibited a weak but distinct eGFP signal in a spatial pattern resembling *β2-tubulin* – absent at the apical germline stem cell region and present in developing cysts downstream. Fluorescence intensity in Draup-A was substantially lower than in AgPMB1 (**Figure 5B**), consistent with the relative abundance of *draupnir* versus *β2-tubulin* transcripts (**Figure 4C**). These observations were confirmed by qRT-PCR quantification of I-*Ppo*I expression from dissected tissues. Autosomal X-shredder strains, regardless of the promoter, displayed robust I-*Ppo*I expression in testes (**Figure 5C**). In contrast, Draup-Y males showed no significant elevation of I-*Ppo*I transcript levels in testes relative to heads, indicating that Y-chromosome context suppresses transcription from the *draupnir* promoter despite its endogenous activity. Lower but still significant I-*Ppo*I expression was also detected in the MAGs of AgPMB1 males. This likely reflects contamination by sperm during tissue dissection, as the *β2-tubulin* promoter is not active in MAGs according to previous data using reporter strains and *in situ* hybridization (Catteruccia, Benton and Crisanti, 2005).

To test whether I-*Ppo*I expression driven by the *draupnir* promoter from either an autosomal or a Y-linked transgene could induce sex ratio bias in offspring, fifty transgenic males from each strain were crossed *en masse* to fifty virgin wild-type females. Eight replicate cages were established per strain and offspring were reared to the pupal stage for sexing. Three replicate G3 wild-type control cages were included in parallel. Compared to G3 crosses (49.5±0.9% males), Draup-A males produced significantly male-biased progeny (82.1±1.5%; Welch’s t-test, p < 0.0001), whereas the offspring sex ratio from Draup-Y crosses (50.1±0.9%) did not differ significantly from G3 (p = 0.60; **Figure 5D**). Sex distortion was further evaluated using females laying eggs individually in single cups, to determine whether mild distortion in a subset of males could be masked by pooled sampling. Male bias was again observed for Draup-A (83.3±1.2%) but not Draup-Y (46.8±1.3%) (**Figure 5D**). Together, these results confirm that while the *draupnir* promoter can drive I-*Ppo*I expression and functional X-shredding from an autosomal locus, the transgene is silenced when integrated on the Y chromosome, consistent with meiotic sex chromosome inactivation (MSCI) acting on Y-linked sequences.

## Discussion

Y chromosome SRDs, or “Y-drives,” offer a compelling route for population suppression of malaria mosquitoes by biasing populations toward males through the targeting of X-bearing gametes. Unlike autosomal SRDs, Y-drives are transmitted exclusively from fathers to sons and are thereby insulated from negative selection in females, enabling efficient and self-sustaining spread once released. Despite their theoretical promise, progress toward implementing Y-linked SRDs in *An. gambiae* has been hindered by the transcriptional silencing of the Y chromosome during male meiosis (Alcalay *et al*., 2021). This phenomenon, known as meiotic sex-chromosome inactivation (MSCI), effectively blocks the expression of canonical meiotic promoters such as *β2-tubulin* when these are linked on the Y chromosome. Here, we identify *draupnir* as the only Y-linked gene in mosquitoes that retains transcriptional activity during meiosis, providing a natural example of transcriptional escape and explore whether this is an intrinsic ability of its flanking 5’ regulatory regions.

Our study first reconstructed the evolutionary origin, genomic organization, and expression of *draupnir*, a multicopy Y-linked gene of *An. gambiae* that encodes a Zip3-like meiotic protein. Phylogenetic analysis indicates that *draupnir* arose through a stepwise duplication process from the ancestral *vilya* homolog, first giving rise to the autosomal paralog *skirnir* early in *Anopheles* evolution (∼100–120 Mya), and later moving to the Y chromosome in the *An. gambiae* complex, where it underwent local amplification. The nearly complete sequence homogenization of *draupnir* copies suggests ongoing concerted evolution within the Y array, potentially mediated by gene conversion or unequal recombination between repeats. The interspersion of *draupnir* copies with *changuu* transposable elements further implies that transposition has contributed to the local spread of the array, echoing the role of repeats in shaping ampliconic domains on mammalian Y chromosomes. Expression analyses confirmed that both *skirnir* and *draupnir* are transcribed in testes, with *draupnir* transcripts peaking during the meiotic stages of spermatogenesis. These results extend earlier findings (Hall *et al*., 2016) by showing that transcription from the Y-linked *draupnir* array persists during the same developmental window in which most other loci are silenced by MSCI. This property places *draupnir* among a rare subset of Y genes that retain transcriptional competence under otherwise repressive chromatin conditions. Notably, while *draupnir* and *skirnir* retain the overall Zip3-like protein fold, both carry an H-D substitution within the RING finger domain that likely alters their E3 SUMO ligase activity relative to the ancestral *vilya*. Whether this modification is functionally consequential or merely a neutral divergence is unclear. The preservation of Zip3-like domain architecture, combined with expression in secondary spermatocytes, suggests that *draupnir* may contribute to meiotic processes or sperm maturation, although its precise role remains unknown.

We hypothesized that the *draupnir* promoter might contain intrinsic features that enable expression from the Y chromsome. However, while the promoter supported meiotic expression and functional X-shredding from an autosomal locus, it failed to drive detectable expression when inserted on the Y chromosome. This demonstrates that meiotic transcription of *draupnir* is not encoded solely in its proximal regulatory sequence. Instead, it likely depends on higher-order features of the Y-linked array, such as copy number, local chromatin environment, repeat organization, or nuclear positioning. These results underscore a major constraint for engineering Y-linked SRDs and suggest that successful strategies will likely require mimicking the native genomic context of expressed Y-linked loci. Potential approaches include multi-copy integration, targeted insertion within permissive Y-linked arrays such as *draupnir*, or incorporation of chromatin elements that promote local transcriptional accessibility. Identifying the features that enable MSCI escape will be essential for advancing Y-linked genetic control strategies. In summary, *draupnir* illustrates how duplication, amplification, and genomic context shape the evolution and function of Y-linked genes. Its transcriptional activity during meiosis provides a valuable model for understanding MSCI escape, while its context dependence highlights a key barrier to engineering Y-linked gene drives in *Anopheles*. Overcoming this barrier will be central to realizing the full potential of Y-linked genetic control strategies.

## Methods

### Identification of *draupnir* homologs in Diptera

Predicted tBLASTn hits of *Ag_vilya, skirnir* and *draupnir* orthologs were curated and gene and protein sequences were downloaded from Vectorbase: *Cx. quinquefasciatus* (CPIJ012237), *Ae. aegypti* (AAEL015042), *An. albimanus* (AALB006177), *An. darlingi* (ADAC002730), *An. minimus* (AMIN006205, AMIN010088), *An. dirus* (ADIR00363, ADIR011231), *An. farauti* (AFAF017148, AFAF004687), *An. atroparvus* (AATE004589, AATE016748), *An. sinensis* (ASIS012546, ASIS022724), *An. melas* (AMEC019848, AMEC011994, AMEC014449), *An. Merus* (AMEM011445, AMEM003621), *An. christyi* (ACHR004360, ACHR000229), *An. quadriannulatus* (AQUA001084, AQUA003841), *An. arabiensis* (AARA009086, AARA005061), *An. coluzzii* (MOPTI, ACMO009001, ACMO006749) and *An.gambiae* (AGAP006749, AGAP013757). In cases where gene annotations exhibited incompleteness or inaccuracies, necessary refinements were applied to ensure accuracy. Amino acid sequences of the *D. melanogaster vilya* (FBgn0283545) and its paralogs, *narya* (FBgn0031018) and *nenya* (FBgn0051053) were downloaded from FlyBase. After screening and sequence polishing, 35 sequences were selected to construct a final phylogenetic tree to determine the evolutionary relationships among the *draupnir*-like orthologs. Phylogenetic analyses were performed using the NGPhylogeny.fr platform (Lemoine *et al*., 2019). Sequences were aligned with MAFFT (Katoh *et al*., 2002) using a gap opening penalty of 1.53 and a gap extension penalty of 0.123. Alignment columns were trimmed with BMGE (Criscuolo and Gribaldo, 2010) using a sliding window of 3, a maximum entropy threshold of 0.5, a gap rate cutoff of 0.5, a minimum block size of 5, and the BLOSUM62 substitution matrix. Phylogenetic trees were inferred with FastTree (Price, Dehal and Arkin, 2010) under the “GTR” model for nucleotide sequences and the “LG” model for amino acid sequences, with gamma-distributed rate variation and branch support estimation. Trees were exported in Newick format and visualized using iTOL (Letunic and Bork, 2021).

### BAC library screening and BAC sequencing

A previously-constructed pooled BAC library made from male genomic DNA of the *An. gambiae* G3 strain (Bernardini *et al*., 2014) composed of approximately 500 primary clones was screened by PCR using *draupnir-*specific primers (**Supplementary Table 1**). These primers were designed to contain *draupnir-*specific SNPs and produced male-specific amplicons in PCR of male and female genomic DNA. Two positive clones were identified, BAC-2C and BAC-5B, which were isolated using the QIAGEN Large-Construct kit (QIAGEN) for and sequenced using Single-Molecule Real-Time (SMRT) PacBio sequencing. BAC sequences were annotated by BLAST analysis using Y chromosome-specific repetitive elements and *draupnir* flanked by 2 kb of putative regulatory sequence from YDB (Hall *et al*., 2016). To confirm Y chromosome linkage of the *draupnir* array *An. gambiae* male and female Illumina whole genomic sequencing data from the G3 strain (Hall *et al*., 2016) were mapped to the PacBio reads using Bowtie (Langmead *et al*., 2009) allowing no mismatches (-v 0) and reporting all alignments (-a). For plotting, coverage for each library was calculated with BEDtools (Quinlan and Hall, 2010) and normalized to library size differences between male and female datasets. Coordinate-based coverage was plotted using custom python scripts.

### Mosquito rearing

Mosquitoes were reared in environmental chambers at 27°C and 80% relative humidity under a 12:12 h light:dark photoperiod. Larvae were reared in plastic trays containing deionized water supplemented with 0.3 g/L of artificial sea salt and fed daily with 5 mL of a 2% (w/v) larval diet, as described by (Damiens *et al*., 2012). Adults were provided *ad libitum* with a 10% sucrose solution containing 1% methylparaben and kept in 30 x 30 x 30 cm plastic-framed cages covered with netting (Bugdorm). Five- to-seven-day old females were blood-fed using a Hemotek membrane feeding system to induce egg production. 48 hrs post-blood feeding, a Petri dish containing a fully hydrated sponge overlaid with filter paper was placed inside the cage to allow oviposition. After hatching, larvae were transferred into the trays containing rearing water. Transgenic individuals were screened for the fluorescent markers using a fluorescent microscope or the Complex Object Parametric Analyzer and Sorter (COPAS; Union Biometrica, Boston, USA).

### RT-PCRs

Total RNA was extracted from dissected male tissues (heads, carcasses, male accessory glands (MAGs) and testes) and female tissues (heads, carcasses and ovaries) using TRIzol reagent (Invitrogen), followed by DNase treatment with the TURBO DNA-free^TM^ kit (Invitrogen). cDNA was synthesized using Superscript III Reverse Transcriptase (Invitrogen) and used for both qualitative and quantitative expression analyses. Qualitative RT-PCR was performed using Phusion^®^ High-Fidelity PCR Master mix (*New England Biolabs*) and primers to amplify *draupnir* and the S7 control gene (**Supplementary Table 1**). Quantitative RT-PCR (qRT-PCR) was performed on cDNA using the iTaq^TM^ Universal SYBR^®^ Green Supermix (Bio-Rad). Expression levels of *draupnir* and I-*PpoI* were normalized to the ribosomal gene *Rpl19* using gene specific primers (**Supplementary Table 1**).

### RNAseq analysis

Stage-resolved RNA-seq data from Taxiarchi et al. (2019), profiling four spermatogenic cell populations (premeiotic, primary spermatocytes, secondary spermatocytes, and postmeiotic cells; NCBI accession PRJEB35368), were used to quantify expression of *vilya*, *skirnir*, *draupnir*, *zpg*, *vasa*, and *β2-tubulin*. Reads from each stage were aligned against a reference comprising the coding sequences of these six genes using Bowtie (Langmead *et al*., 2009) with parameters -v 0 (no mismatches permitted) and -a (reporting all valid alignments). SAM files were converted to sorted BAM files using SAMtools (Li *et al*., 2009). Read counts were extracted from BAM files and normalized to transcripts per million (TPM). Alignments were additionally performed using HISAT2 (Kim *et al*., 2019) with the -a flag, and the fraction of uniquely versus multi-mapping reads was assessed to confirm the consistency of expression estimates; Bowtie-derived results are presented as they were concordant between the two methods.

### *K-mers* analysis

To distinguish paralog-specific expression between the highly similar skirnir (autosomal) and draupnir (Y-linked, ten copies) sequences, we used a k-mer-based approach. Using Jellyfish (Marçais and Kingsford, 2011), we generated two sets of gene-specific 25-*mer* sequences: one unique to *skirnir* and one unique to *draupnir*. Staged RNA-seq data from Taxiarchi et al. (2019), which profiled four spermatogenic cell populations (premeiotic, primary spermatocytes, secondary spermatocytes, and postmeiotic cells; NCBI accession PRJEB35368, three replicates per stage), were used to extract and count matching *k-mers* using the Jellyfish (Marçais and Kingsford, 2011) *k-mer* counting tool. RNA *k-mer* counts were normalized by sample size. Mean *k-mer* counts per spermatogenic stage were calculated and visualized using custom Python scripts.

### Construction of transformation plasmids and transgenic strains

The putative *draupnir* 5’ regulatory region was amplified from male genomic DNA using primers designed around Y-chromosome specific SNPs (**Supplementary Table 1**) and was cloned upstream of an I-*Ppo*I X-shredder cassette based on the destabilized I-PpoI*^124L^* variant fused to eGFP with the F2A ribosome skipping site and terminated by the *β2-tubulin* 3’ UTR (Galizi *et al*., 2014). This construct contained an attB site for site-specific integration and a promoterless eCFP fluorescence marker that replaces the RFP marker present in the recipient attP site upon integration. Embryos from the AttP-RFP-H2 and Y-attP strains were microinjected with a mixture containing 0.05 mg/mL of the transformation plasmids and 0.4 mg/mL of a helper plasmid encoding a *vasa*-driven *φC31* integrase. G_0_ adults displaying transient eCFP fluorescence at larval stages were crossed to wild-type mosquitoes to F1 offspring that were screened for the fluorescent marker. Transgenic F1 were back-crossed to wild types to establish Draup-A and Draup-Y transgenic strains.

### Confocal microscopy

Testes were dissected from males of the wild-type G3, AgPMB1, Draup-A and Draup-Y lines and fixed in methanol-free 16% formaldehyde (Pierce) in PBS for 30 minutes. Samples were then washed three times for 15 minutes each in PBS containing 0.1% Tween-20. Fixed testes were mounted on glass slides in Vectashield Antifade Mounting Medium with DAPI (Vectorlabs) and covered with coverslips. Images were acquired using a Zeiss LSM 510 laser scanning confocal microscope with a 20x objective using the eGFP channel. The exposure was increased up to 50-fold to detect fluorescence intensity in Draup-A testes, whereas no fluorescence signal was observed in Draup-Y testes under the same settings.

### Sex-ratio distortion assays

Sex distortion assays for the Draup-A and Draup-Y transgenic lines and G3 as a control were conducted in small cages (30 x 30 x 30 cm) by crossing 50 transgenic males with 50 wild-type females. Mosquitoes were allowed to mate for 5 days and subsequently blood-fed to allow females to lay eggs. The progeny were screened to assess sex bias at the pupal stage. In parallel, the assay was repeated using the same number of males and females, but females were transferred individually to separate cups.

## Author Contributions

RDA, AC and PAP conceived and designed the study. RDA and AC generated transgenic constructs and strains and performed all mosquito experiments. ESY, FK and AS performed the orthology and bioinformatics analyses. SDM, AT helped in mosquito experiments. RG, NW and AS provided key insights and support. PAP supervised the project, secured funding. RDA, ESY and PAP wrote the manuscript, with input from all authors. All authors read and approved the final manuscript.

## Acknowledgements

We thank Francesco Papa for the initial bioinformatic analysis of *draupnir.* We are thankful to Nora Besansky, Jake Tu, Igor Sharakhov, Adam Phillippy, Changde Cheng, Matt Hahn and other members of the *Anopheles* Y consortium, which initiated the work on *draupnir.* We thank Dr. Jaroslaw Krzywinski and Roberta Spaccapelo for their generous and continuous support.

## Ethics

Mosquito experiments were conducted at the Department of Experimental Medicine, University of Perugia and Polo GGB (Terni), which hold Arthropod Containment Level 2 (ACL2) certification. Containment protocols and risk assessment measures for genetically modified arthropods were approved by the Italian Ministry of the Environment (MATTM) and the Italian Institute for Environmental Protection and Research (ISPRA). Research involving the contained use of genetically modified microorganisms is authorized by the Italian Ministry of Health under Legislative Decree 206/2001.

---

**Supplementary Table 1:**
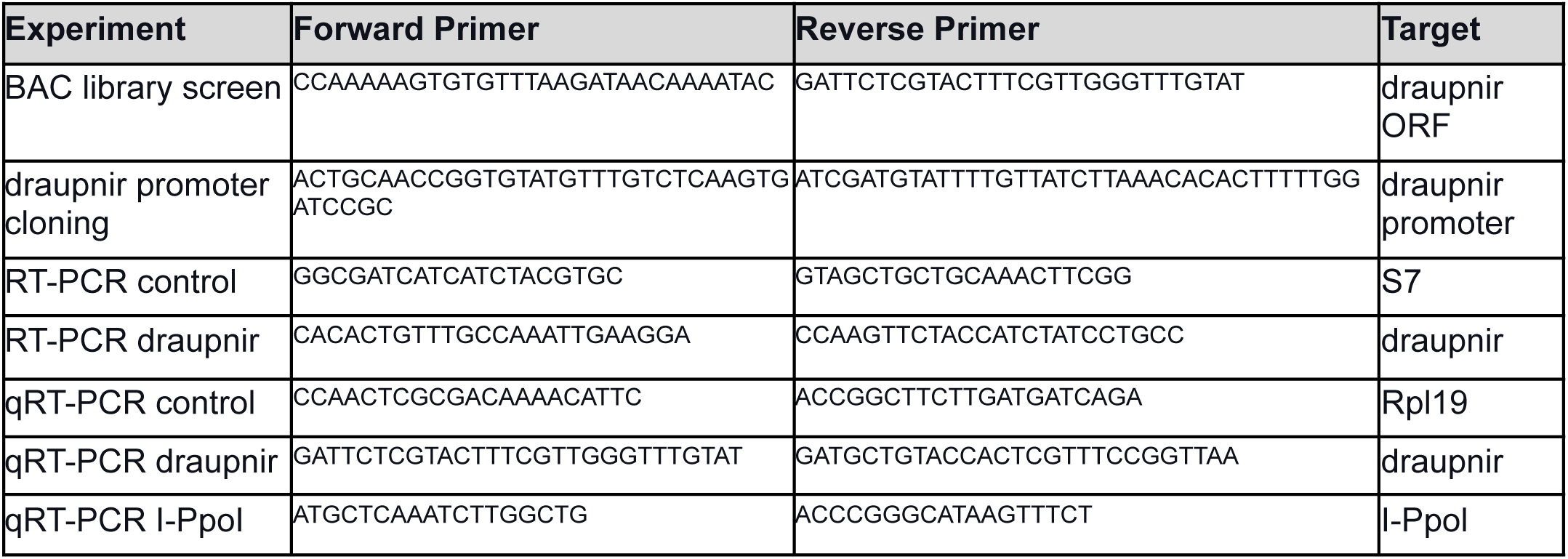
Primers used in this study.

**Supplementary Figure S1:**
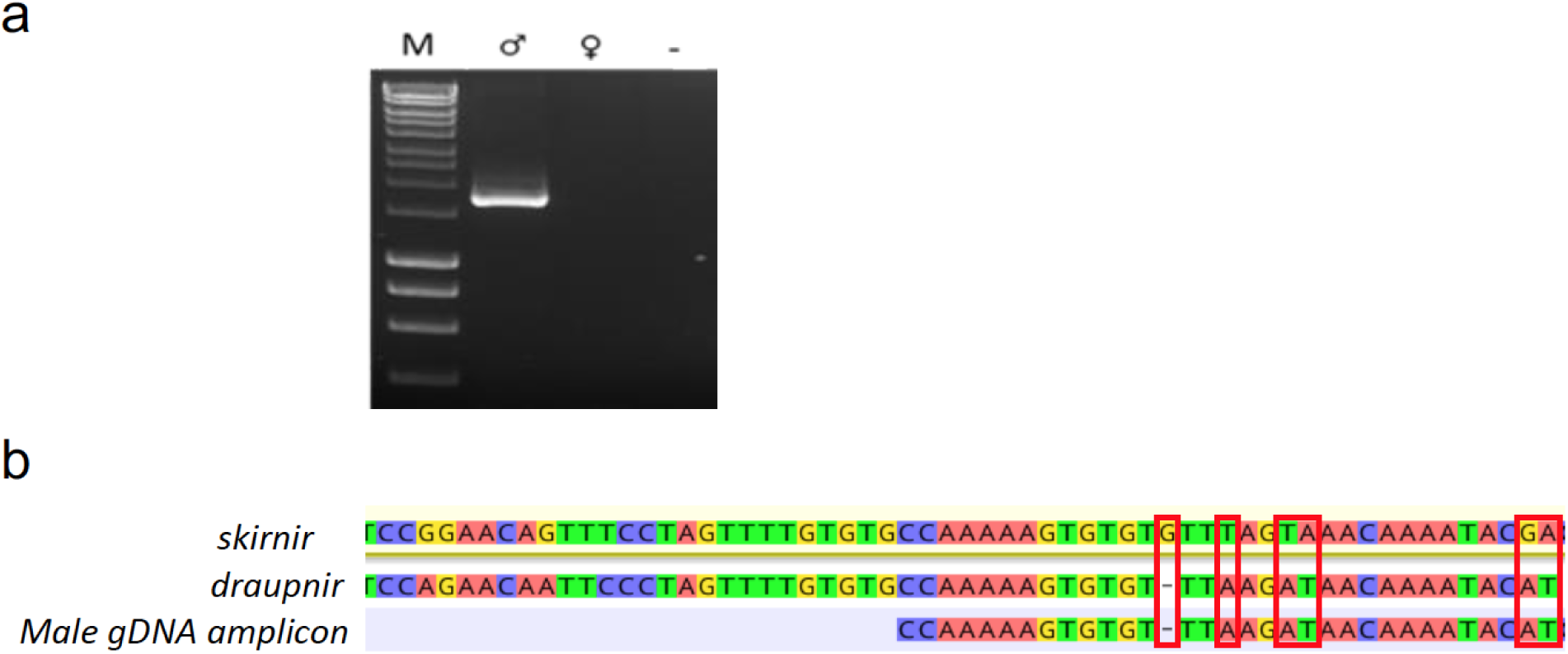
(a) PCR amplification from male and female genomic DNA using primers spanning the *draupnir* 5’ regulatory region. (b) Detection of *draupnir*-specific SNPs based on sequencing of male genomic DNA amplicons confirm amplification of the Y-linked *draupnir*.

